# Digging behavior discrimination test to probe burrowing and exploratory digging in male and female mice

**DOI:** 10.1101/2020.12.29.424478

**Authors:** Heather L. Pond, Abigail T. Heller, Brian M. Gural, Olivia P. McKissick, Molly K. Wilkinson, M. Chiara Manzini

**Affiliations:** Department of Pharmacology and Physiology and Integrative Systems Biology, The George Washington University School of Medicine and Health Sciences, Washington, DC 20037, USA; Child Health Institute of New Jersey and Department of Neuroscience and Cell Biology, Rutgers Robert Wood Johnson Medical School, New Brunswick, NJ 08901, USA

**Keywords:** repetitive behaviors, behavioral analysis, burrowing, digging

## Abstract

Digging behavior is often used to test motor function and repetitive behaviors in mice. Different digging paradigms have been developed for behaviors related to anxiety and compulsion in mouse lines generated to recapitulate genetic mutations leading to psychiatric and neurological disorders. However, the interpretation of these tests has been confounded by the difficulty of determining the motivation behind digging in mice. Digging is a naturalistic mouse behavior, that can be focused toward different goals, i.e. foraging for food, burrowing for shelter, burying objects, or even for recreation as has been shown for dogs, ferrets, and human children. However, the interpretation of results from current testing protocols assumes the motivation behind the behavior often concluding that increased digging is a repetitive or compulsive behavior. We asked whether providing a choice between different types of digging activities would increase sensitivity to assess digging motivation. Here, we present a test to distinguish between burrowing and exploratory digging in mice. We found that mice prefer burrowing when the option is available. When food restriction was used to promote a switch from burrowing to exploration, males readily switched from burrowing to digging outside, while females did not. In addition, when we tested a model of intellectual disability and autism spectrum disorder that had shown inconsistent results in the marble burying test, the *Cc2d1a* conditional knock-out mouse, we found greatly reduced burrowing only in males. Our findings indicate that digging is a nuanced motivated behavior and suggest that male and female rodents may perform it differently.

**Significance Statement:** Digging behavior is commonly assessed in mice to study features of neurodevelopmental, psychiatric and neurological disorder. However, existing digging assays fail to discriminate between types of digging complicating data interpretation. Here we present a modified digging behavior discrimination task that can produce sensitive results in 30 minutes with easy to gather measures, making it accessible to wide variety of labs and experimental paradigms.

## Introduction

The innate digging and burrowing behaviors displayed by house mouse (*Mus musculus*) strains commonly used in the laboratory are valuable indicators of well-being and motor function (Dudek et al., 1983; Latham & Mason, 2004), and are used to test pain, stress, and features of neurological and psychiatric conditions such as anxiety, Autism Spectrum Disorder (ASD), and Obsessive-Compulsive Disorder (OCD) (Deacon, 2006b; Deacon et al., 2001; Jirkof, 2014; de Brouwer et al., 2019).

Mice dig for a number of reasons; to avoid noxious stimuli or predators, to seek food, to build shelter for safely raising their young, and possibly for recreation (Arakawa et al., 2007; Latham & Mason, 2004; Powell & Banks, 2004; Sluyter et al., 1996). Deep bedding will induce a mouse to dig into the substrate (Deacon, 2006b), but the motivation behind this behavior remains uncertain. Increased digging is often interpreted as a repetitive behavior due to anxiety-like and compulsive-like responses (Broekkamp et al., 1986; Thomas et al., 2009). However, a compulsive behavior is defined as excessive and divorced from the consummatory process, i.e. not leading to pleasure or reward (American Psychiatric Association, 2013; Luigjes et al., 2019). Defining whether an activity is pleasurable or excessive is difficult to assess in mice since the motivation for digging is often unknown. Thus, free digging is also used as a measure of a more generic exploratory drive instead (de Brouwer et al., 2019).

One of the most commonly used digging tests is the marble burying test where marbles are placed on the digging surface and the act of embedding an object in the substrate is studied (Broekkamp et al., 1986). The validity of interpreting marble burying as a sign of anxiety-like or compulsive-like behavior has been challenged in multiple studies revealing a need to define the motivation behind digging (Bruins Slot et al., 2008; Gyertyán, 1995; Hayashi et al., 2010). It remains unclear whether mice actively interact with the marbles as novel or aversive objects or whether burying (and unburying) is simply a side effect of vigorous digging in the vicinity (Gyertyán, 1995; Thomas et al., 2009).

Burrowing, the act of digging for shelter, has been studied in multiple species of rodents and defined as a mandatory behavioral need for laboratory mice by Sherwin et al., (2004). A mandatory behavioral need is a natural behavior whose functional consequences are clearly important to the animal who is strongly motivated to perform it, as observed in previous burrowing studies (Deacon, 2006b; Jirkof et al., 2010). While studying burrow building requires a large apparatus, the act of burrowing can be tested in laboratory settings by providing a tube filled with bedding that mice can clear. This protocol was developed by Deacon (2006a) exploring both interaction with food pellets or other non-food related substrates and allowing the mice to burrow for multiple hours.

To develop measures to discern the individual motivation for digging behavior we combined burrowing and free digging assays in a single paradigm. Our approach, the digging behavior discrimination (DBD) task, applies the burrowing method described by Deacon (2006a), truncated to 30 minutes and modified to include measurement of free digging. This assay was tested in both male and female mice to define its sensitivity to assess changes in digging during an environmental challenge (food restriction) and in a mouse strain recapitulating loss of a human gene leading to intellectual disability (ID) and ASD. We identified multiple differences between male and female mice under food restriction and with the ID/ASD mutation. While previous studies have not reported any sex differences in digging behavior, the DBD test shows there are differences in digging motivation between males and females and allows for clear differentiation between exploratory digging/foraging and burrowing.

## 2. Materials and Methods

### 2.1 Animals

All animal care and use were in accordance with institutional guidelines and approved by the Institutional Animal Care and Use Committee of The George Washington University and Rutgers University. All animals were maintained in group housing (5 animals/cage maximum) in ventilated cages from Tecniplast USA (West Chester, PA) with corncob bedding (Bed-o’cobs, Anderson, Maumee, OH) at 20-26 °C and 30-70% humidity on a 12-hr light/dark cycle. Enrichment was provided as shredding nestlets. Cages were changed every 2 weeks by designated facility staff. C57BL/6N male and female mice (Males M1: N=11; M2 N=13; Females N=10) were purchased from Charles River Laboratories (Frederick, MD; RRID: IMSR_CRL:27) and Taconic (Albany, NY; RRID:IMSR_TAC:b6) and randomly assigned to the different cohorts. Mice were acclimated in house for at least eight weeks to account for differences among suppliers and tested at around 4-5 months of age. The *Cc2d1a* conditional knock-out (cKO) mouse line was generated by crossing *Cc2d1a*-flx mice (Oaks et al., 2017, RRID: MGI:5449582) with a CaMKIIa-cre mouse line driving Cre recombinase expression under the *CaMKIIa* promoter (Jackson Laboratories, RRID:IMSR_JAX:005359) (Tsien et al., 1996). All experimental animals (Control M N=8; cKO M n=8; control F N=10; cKO F n=10) are fully backcrossed on a C57BL/6N background (RRID: IMSR_CRL:27) for at least 6 generations. Genotyping was performed via polymerase chain reaction (PCR) amplification and primers are available upon request. Experimental animal numbers were chosen upon power analysis to detect at least a 30% difference in performance with 90% confidence.

### 2.2 Burrowing and exploratory digging discrimination

Testing was performed starting at 10 am and animals were brought to the testing room in their home cages 1-2 hours prior the beginning of testing. Illumination in the room was provided only through a red light and a white noise machine was used to dampen possible ambient noises. The test was performed in a clear plastic box 40X24X31.75cm (X-Large Kritter Keeper, Lee’s Aquarium and Pet Products, San Marcos, CA). The box was filled with 5cm of Bed-o’cobs bedding to provide ample digging substrate. A “burrow” consisting of a yellow transparent plastic tube (10cm length, 5cm diameter) filled with 17g of white Carefresh paper bedding (Healthy Pet, Ferndale, WA) was placed in a corner of the testing arena by gently pressing into the substrate to prevent rolling (**Fig. 1A**). To familiarize the mice with the burrow and eliminate the confound of a novel object, a burrowing tube filled with the paper bedding was placed in the home cage of the group-housed test mice the night before testing. Testing was only performed if the tube was empty by the following day. If not, one more night of habituation was granted to assure the mice were able to demonstrate burrowing behavior. On testing day, each mouse was placed in the test apparatus and movement was tracked for 30mins using AnyMaze software (Stoelting, Wood Dale, IL) between two testing zones: the burrow area and the rest of the box or “exploration area”. Multiple automated testing measures were collected, including time in burrow area, time in exploration area, number of entries in the burrow area, average time per visit, average speed in apparatus, and distance traveled in apparatus. Latency to start removing material from the burrow, time spent burrowing or digging, and time to empty the burrow were timed manually from the videos by two independent raters masked to genotype. Digging in the free area was defined as vigorous digging with spread hind limbs and coordinated use of the forefeet to move substrate backwards beneath the body or by the sides as previously described (Layne & Ehrhart, 1970; Webster et al., 1981). The weight of the bedding left in the burrow was weighed at the end of the test.

**Figure 1.**
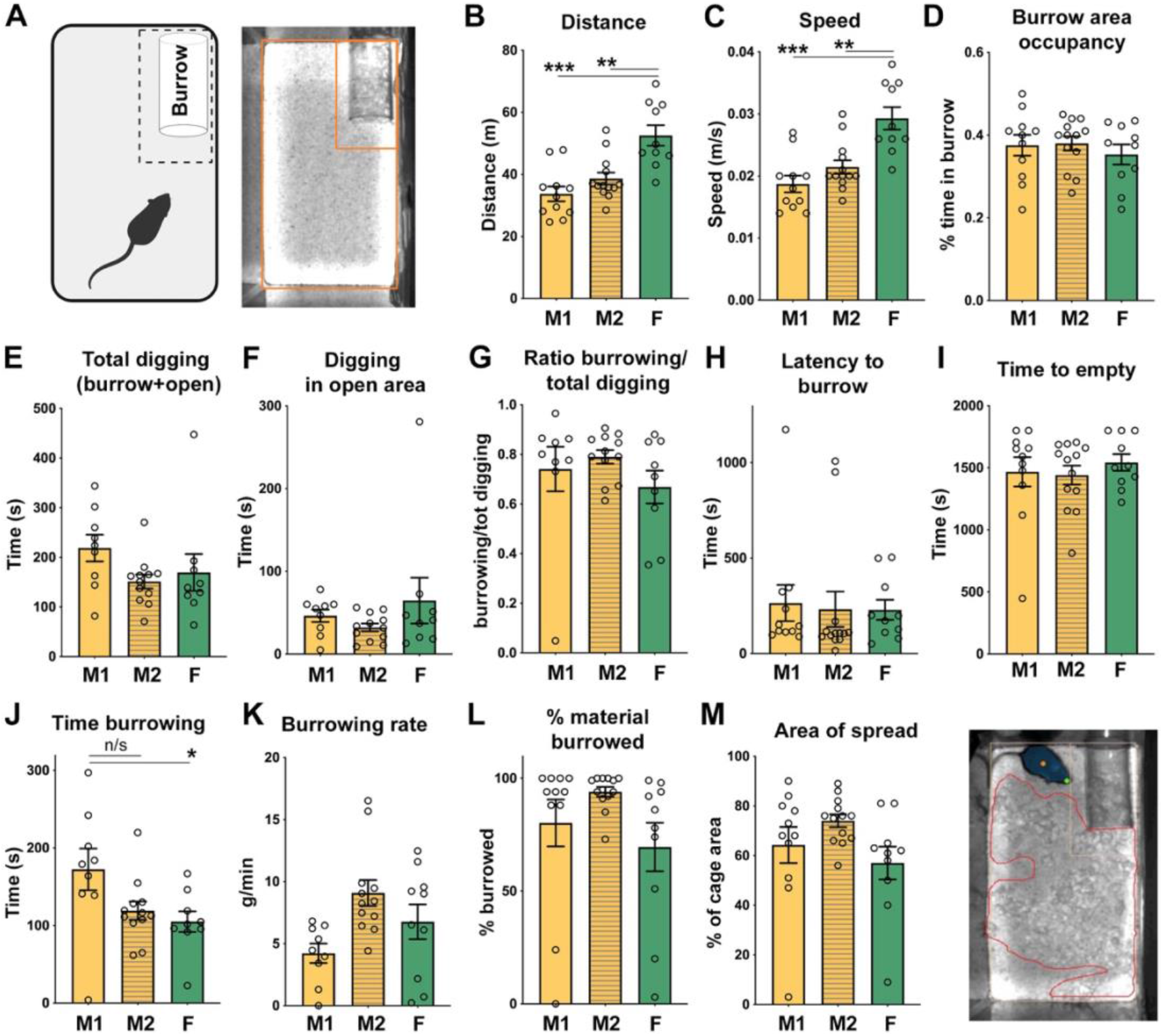
Stability of performance on Digging Behavior Discrimination test. Three cohorts of mice (M1: N=11; M2 N=13; F: N=10) were tested independently to assess stability of burrowing and digging performance and define possible sex differences. (***A***) Test chamber set up and schematic of digging and burrowing zones. (***B-C***) Males (M1, M2) covered similar distances at equal speed, while females showed increased motor activity (**B**) and speed (**C**). (***D-F.***) Different cohorts showed mostly consistent digging performance inside and outside the burrow area. Occupancy in the burrow area (**D.**), total time spent digging in both areas (**E**.) and outside (**F.**) were similar. (**G.**) Most of the digging time was spent burrowing. (**H-I**.) Latency to burrow and time to empty the burrow were also similar in all cohorts. (**J.**) Time spent in direct interaction with substrate in the burrow was variable with females significantly lower from the M1 group, but not M2. (**K.**) The burrowing rates, grams of burrow substrate removed per minute, were not significantly different, but M1 trended towards a slower rate. (**L.**) Overall, most animals efficiently removed the burrowing substrate from the tube by the end of the test. (**M.**) The substrate was then distributed over the area of the cage (example of spread soft bedding outlined in red on the right). Values are presented as means ± *SEM*. Symbols are individual mouse data points. \**p* < 0.05, **<0.01, or ***<0.001 following multiple comparison tests. All unmarked comparisons were not significant. Additional statistical information is reported in Suppl. Table 1.

### 2.3 Food restriction

Food restriction was performed following baseline testing by gradually decreasing daily food rations from 5g to 1g in a group-housed setting until each animal lost up to 15% of its initial weight in the span of 2 weeks. Animals were weighed daily and would be isolated only when one animal was lagging behind in its weight loss and found to be consuming more food than the others. This was done to reduce the possible confound of isolating all animals for food restriction. Animals were tested again as described above without the overnight habituation period and returned to *ad libitum* diet for 2 weeks with their usual group.

### 2.4 Corticosterone testing

Submandibular blood collection method was used to obtain samples under isofluorane anesthesia. A sterile, stainless steel lancet (MEDIpoint Inc., Mineola, NY) was used to pierce slightly behind the mandible to collect a 100uL blood sample in an EDTA microtainer blood collection tubes (BD Diagnostics, Franklin Lakes, NJ). The collection tubes were then spun at 2000 rpm for 10 minutes to separate the plasma from the blood sample. Corticosterone levels in the plasma were measured using the Detect X^®^ Corticosterone Enzyme Immunoassay Kit (Arbor Assays, Ann Arbor, MI) on a Varioskan LUX multimode microplate reader (Thermo Scientific, Waltham, MA), following the manufacturer’s instructions.

### 2.5 Statistical analysis

All datasets were tested for normality using the Shapiro-Wilk test and appropriate statistical test were applied. One-way ANOVA or the Kruskal-Wallis test were used for baseline cohort measures with respectively Tukey’s or Dunn’s multiple comparison tests. Two-way ANOVA was used for the food restriction studies (with repeated measures) and to analyze the *Cc2d1a* cKO cohorts to determine the effect of treatment (food restriction or genotype) and sex with Tukey’s multiple comparison test. One-tailed Pearson correlation was performed to determine correlation whenever multiple comparisons were not needed. Repeated measure correlation was performed using the Rmcorr package in R to compare variable in the same animals during food restriction studies (Bakdash & Marusich, 2017)

## 3 Results

### 3.1 Test design for Digging Discrimination test

We sought to develop a novel paradigm to discern the motivation behind different digging behaviors. The test design was based on a combination of existing tests, a burrowing test (Deacon, 2006a), and free digging (Deacon, 2006b). We chose a box larger than the home cage and similar to the one used for marble burying and free digging tests to provide space for movement and exploration. The testing arena was filled with a thick (5 cm) layer of corncob bedding, the same as bedding used in their home cage to provide a familiar digging substrate. The “burrow” consisted of a plastic tube as used for burrowing in Deacon (Deacon, 2006a). While the Deacon test packed the tube with food pellets or pea shingles requiring at least 3 hours of testing per animal, we used soft bedding allowing for faster testing times since we found that mice would not burrow readily with higher packing densities or heavier materials. The type of bedding and packing weight of the tube (17g) was determined by testing different packing densities and identifying the optimal amount of bedding that could be completely removed in less than 30 minutes by a wild type mouse. To remove the confound of interacting with a novel object and pre-train the animal for bedding removal, habituation to the tube filled with bedding was performed in the home cage the night before testing.

At the beginning of the test, the burrow tube was placed in a corner of the testing apparatus. For automated video tracking the area surrounding the tube was outlined as the burrowing area to also capture activity close to the tube and the remaining area was used to monitor movement and exploratory digging activity (**Fig. 1A**). Each mouse was placed in the corner opposite to the burrow and multiple parameters were tracked for 30 minutes via either automated video tracking or video analysis by independent raters. Automated parameters included basal activity levels such as total distance traveled and speed, and occupancy of the burrowing and free digging areas. Interaction with the burrow was quantified by measuring the latency to enter the burrow area, quantifying the time spent interacting with the substrate in the burrow (digging or pushing bedding outside), and by manually recording the time to empty the burrow. Digging in the open area was also quantified limiting the analysis to vigorous digging as defined in the Methods. At the end of the test, the soft bedding filling remaining in the burrow was weighed to determine the percent of weight removed.

### 3.2 Male and female performance in the DBD test

Performance, test stability, and sex as a biological variable were assessed by testing two separate age-matched cohorts of C56BL/6N males and one cohort of females (**Fig. 1**. M1: N=11; M2 N=13; F: N=10). Two independent groups of male mice were run at different times to test the stability of the testing conditions and one group of females was run to test for sex as a biological variable. While females showed increased distance covered in the arena (**Fig. 1B**, M1:33.7±2.4 m, M2:38.7±1.9 m, F: 52.6±3.3 m; M1/F p<0.0001, M2/F p=0.0013) and speed (**Fig. 1C**, M1: 0.019±0.001 m/s, M2: 0.021±0.001 m/s, F:0.029±0.002 m/s; M1/F p<0.0001, M2/F p=0.0011), burrowing and exploratory digging performance was comparable among male and female cohorts. Automated tracking of area occupancy indicated that all groups spent around on third of their time in the burrowing area (**Fig.1D**, M1: 0.375±0.025, M2: 0.380±0.017, F: 0.353±0.024, for all non-significant statistics see **Suppl. Table 1**), but careful analysis of digging parameters showed that the automated numbers did not reflect where the mice chose to dig. Total digging activity combining digging in the burrow or in the open area was comparable between males and females (**Fig.1E**, M1: 218.8±26.9 s, M2: 151.3±14.15 s, F: 169.6±36.9 s), but only a small amount of time was spent digging in the open area (**Fig.1F**, M1: 46.3±7.5 s, 32.1±4.7 s, 64.6±27.2). By calculating how much of total digging time was spent burrowing, we found that mice spent more than two thirds of their digging time in the burrow suggesting that burrowing is preferred (**Fig.1G**, burrowing/total digging time M1: 0.74 ±0.09, M2: 0.79±0.03, F: 0.67±0.07). Males and females showed similar latency to interact with the substrate in the burrow (**Fig.1H**, M1:264.5±94.8 s, M2: 232.3±92.5, F: 228.9±52.3). Most animals were able to completely empty the burrow tube within the allotted 30 minutes (1800 s) (**Fig.1I, Suppl. Table 1**).

Two independent raters visually analyzed the videos for burrowing and exploratory digging. Time spent burrowing was significantly reduced in females when compared to M1, but not M2, nor M1 and M2 were statistically significant from each other (**Fig. 1J**, M1: 172.4±26.9 s, M2: 119.2±11.8 s, F: 105.0±13.4 s; M1/F p=0.04, **Suppl. Table 1**). Since most of the M1 cohort had emptied the burrow like the others, we calculated the burrowing rate, i.e. how much bedding in weight was removed per minute of digging, and found that M1 males were significantly slower burrowers than M2 males (**Fig. 1K**, M1: 4.22±0.78 g/min, M2: 9.08±1.05 g/min, F: 6.76±1.40 g/min; M1/M2 p=0.012). Burrowing activity was measured at the end of the test and determining what percentage of the substrate had been removed. Though averages ranged between 94.0±2.2% for the M2 male cohort and 69.5±10.7% for females, no significant differences were observed in burrowing performance (**Fig. 1L, Suppl. Table 1**).

In addition, we noted consistent thin spreading of the soft bedding removed from the burrow on the surface of the cage. Soft bedding was pushed outside of the tube and often methodically distributed around the exploratory area of the arena by spreading it with the nose or front paws in a flicking or wading motion. While the flicking motion was not quantified as we could not determine whether the mice were interacting with the soft bedding or the corncob, we measured how much of the exploration area was covered by soft bedding at the end of the test as a measure of spreading behavior (**Fig. 1M**). There was no significant difference between males and females. The spreading measure showed a positive correlation to the amount burrowed for mice in groups M1 and F (Pearson r M1: r=0.83, p=0.0028; M2: r=0.31, p=0.164; F: r=0.79, p=0.0053) suggesting that the animals may consistently spread the material removed from the burrow. Overall, we found that the 30-minute test was sufficient to completely empty the burrow and dispose of the removed material and to discriminate digging within the burrow and exploratory digging in the outside area. Multiple additional digging and burrowing parameters such as burrowing rate and the ratio of time spent in different digging activities could be collected.

### 3.3 Digging discrimination with food deprivation

To test the sensitivity of the test, we asked if food restriction would change digging preference and elicit a shift between burrowing and exploratory digging/foraging outside the burrow. We performed the digging discrimination test following a food restriction protocol leading to 10-15% weight loss and after *ad libitum* feeding was restored for 2 weeks (**Fig. 2A**). Males from cohort M2 and female mice were used. During the food restriction condition three mice of each sex escaped in the middle of the trial as soon as they emptied the burrow and were excluded from the analysis. Even if these mice completed the test following *ad libitum* feeding, results from these animals were excluded from the final analysis in order to only include animals who completed all three trials. Data on all parameters measured and statistical analyses is reported in **Suppl. Table 2**. Information on the three animals per sex that were excluded has been provided as **Suppl. Table 3** showing that while they appeared more active at baseline, burrowing activity was consistent with the rest of the group.

**Figure 2.**
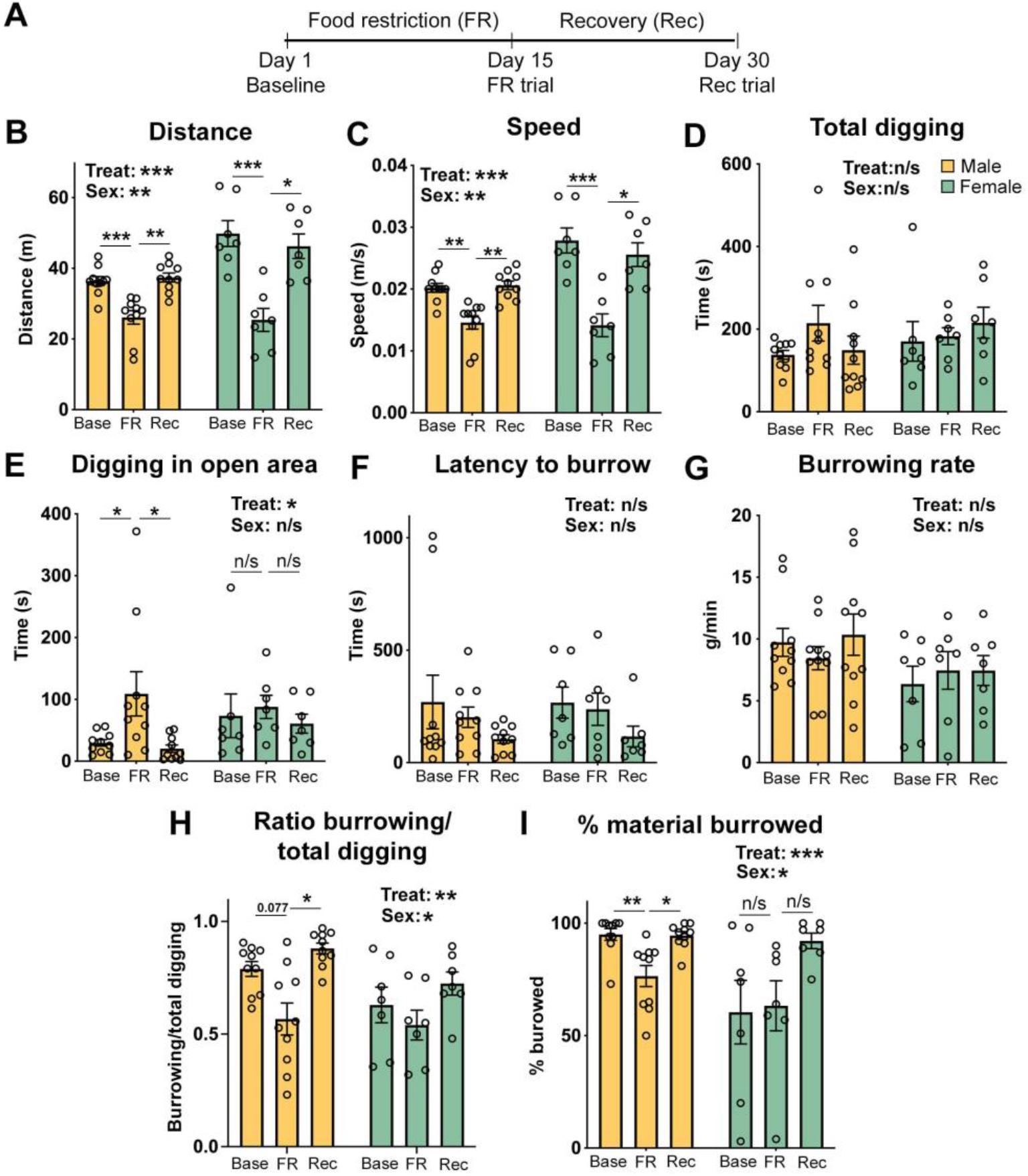
Male and female mice show different free digging and burrowing performance after a food restriction challenge. **(A.)** Two cohorts of mice (Male: N = 10; Female: N = 7) were assessed using the DBD test at baseline (Base), during food restriction (FR), and once recovered from food restriction (Rec). **(B – C.)** Female mice covered more distance (**B.)** at a faster pace (**C.**) than male counterparts at baseline and showed a more prominent drop to male-like levels of activity during FR. Both sexes recovered to baseline levels. (**D.**) Total digging activity remained similar. (**E.**) Males spent significantly more time digging during FR, while females maintained constant digging performance. (**F-G.**) Latency to burrow and burrowing rate did not change significantly during FR or recovery. (**H-I**.) Analysis of the ratio between burrowing and total digging and percentage of material removed from the burrow revealed differences in response between males and females. Females engage in limited burrowing at baseline and FR, but increase during recovery, whereas males burrow substantially at baseline, reduce during FR, and return to baseline performance during recovery. Values are means ± *SEM*. Symbols are individual mouse data points. \**p* < 0.05, **<0.01, or ***<0.001 following multiple comparison tests. All unmarked comparisons were not significant. Additional statistical information is reported in Suppl. Table 2.

Mice of both sexes covered less distance at lower speed after food restriction and returned to baseline during recovery showing a strong effect of the treatment (**Fig. 2B-C**). In addition, a larger effect was noted in females who displayed a larger reduction in mobility than males (**Fig. 2B-C**, Baseline=Base, Food Restriction=FR, Recovery=Rec. Distance. Base: M, 36.4±1.3m; F, 49.9±3.7 m. FR: M, 26.1±1.9 m; F, 25.4±3.2 m. Rec: M, 37.4±1.3 m; F, 46.3±3.5 m. Speed. Base: M, 0.020±0.001 m/s; F, 0.028±0.002 m/s. FR: M, 0.015±0.001 m/s; F, 0.014±0.02 m/s. Rec: M, 0.021±0.001 m/s; F, 0.026±0.002 m/s. 2-way ANOVA: treatment p= <0.0001 for both distance and speed, sex p=0.0075 for distance, sex p=0.0059 for speed, sex X treatment p=0.0074 for distance, p=0.0075 for speed). Both males and females decreased their total time in the burrow area spending more time in the open space (**Suppl. Table 2**). Despite the observed reduction in total mobility after food restriction, total digging time showed no significant changes with a trend for increased digging during food restriction only in males (**Fig.2D**, Base: M, 137.8±10.6 s; F, 170.6±48.0 s. FR: M, 214.4±43.3 s; F: 182.9±20.5 s. Rec: M, 149.2±33.9 s; F, 215.6±37.51 s. Statistics in **Suppl. Table 2**). When exploratory digging was considered alone, it appeared that the increased total digging trend in males was driven by a 3.6-fold increase in time spent digging in the open area following food restriction and no such differences were observed in females indicating a male-specific response (**Fig.2E**, Base: M, 29.8±5.3 s; F, 73.50±35.4 s. FR: M, 109.0±35.7 s, F, 87.9±18.8 s. Rec: M, 20.3±6.2 s; F, 60.7±15.3 s. M-Base/M-FR p=0.032, M-FR/M-Rec p=0.015, other statistics and ANOVA results in **Suppl. Table 2**).

Digging in the burrow was affected more moderately. There was no significant difference and no effect of treatment or sex in the latency to burrow, or burrowing rates, though latencies trended towards faster times with every repetition of the test (**Fig. 2F-G, Suppl. Table 2**). When the ratio between burrowing and total digging activity was calculated significant effects of both treatment and sex emerged showing a possible reduction in burrowing in males and no response to food restriction in females (**Fig. 2H**, Base:M, 0.79±0.03; F 0.63±0.08. FR:M, 0.57±0.07; F, 0.54±0.07. Rec: M, 0.88±0.02; F, 0.72±0.05. M-Base/M-FR p=0.071, M-FR/M-Rec p=0.0014, 2-way ANOVA: treatment p=0.0012, sex p=0.025, sex X treatment p=0.410). The percentage of material burrowed parameter was a better readout for this sex-specific response showing a small significant reduction in material burrowed after food restriction in males (**Fig.2I**, Base: 95.0±2.7%;FR: 76.5±4.7%; Rec: 94.5±1.8%. Base/FR p=0.0017, FR/Rec p=0.024, 2-way ANOVA: treatment p=0.0007). Females were not affected by food restriction revealing an effect of sex on performance and an interaction between sex and treatment (**Fig.2I**, Base: 60.4±14.2%; FR: 63.3±11.1%; Rec: 92.1±3.5%, 2-way ANOVA: sex p=0.041, sex X treatment p=0.014). Interestingly, only on their third test trial after recovery from food restriction females burrowed as efficiently as males (**Fig. 2I**). These results show that the DBD test was sensitive in showing a switch from burrowing to digging in the open space that surprisingly was specific to males. In addition, among all the measures of burrowing the percentage of material removed from the burrow was able to reveal small significant differences in burrowing efficiency.

Since corticosterone (CORT) levels are elevated by food restriction (Guarnieri et al., 2012; Pankevich et al., 2010), we wondered whether they would correlate with digging performance. CORT levels were measured by ELISA during the baseline testing showing that females had higher baseline CORT levels than males as previously observed (Kitay, 1961; Laviola et al., 2002) (**Fig. 3A, Suppl. Table 1**). After food restriction, males followed the expected pattern with an increase in CORT levels and returned back to baseline with *ad libitum* feeding (**Fig. 3B** and **Suppl. Table 2**, Base: 167.5±12.1 ng/μl; FR: 401.4±50.0 ng/μl; Rec: 206.0±13.1 ng/μl. Base/FR p=0.005, FR/Rec p=0.006). Females showed a smaller but significant increase following food restriction, but levels remained elevated in the recovery trial (**Fig.3B**, Base: 336.7±32.1 ng/μl; FR: 480.1±39.3 ng/μl; Rec: 430.9±52.5 ng/μl. Base/FR p=0.009, FR/Rec p=0.765). We performed repeated measures correlation analysis between CORT levels and digging time or percentage of material burrowed for males and females. Males showed a strong positive correlation between CORT and digging (r=0.63, 95% CI [0.24, 0.84], p=0.0024) and strong negative correlation between CORT and material burrowed (r=−0.73, 95% CI [−0.89, −0.42], p=0.00015). Females showed no significant correlation for either digging variable (Digging: r=−0.27, 95% CI [−0.72, 0.33], p=0.325; Material burrowed: r=0.14, 95% CI [−0.44, 0.64], p=0.609). While we cannot conclude that corticosterone is linked to the behavioral changes in males, these results support the finding that males and females show differentially change their digging behavior following food restriction.

**Figure 3.**
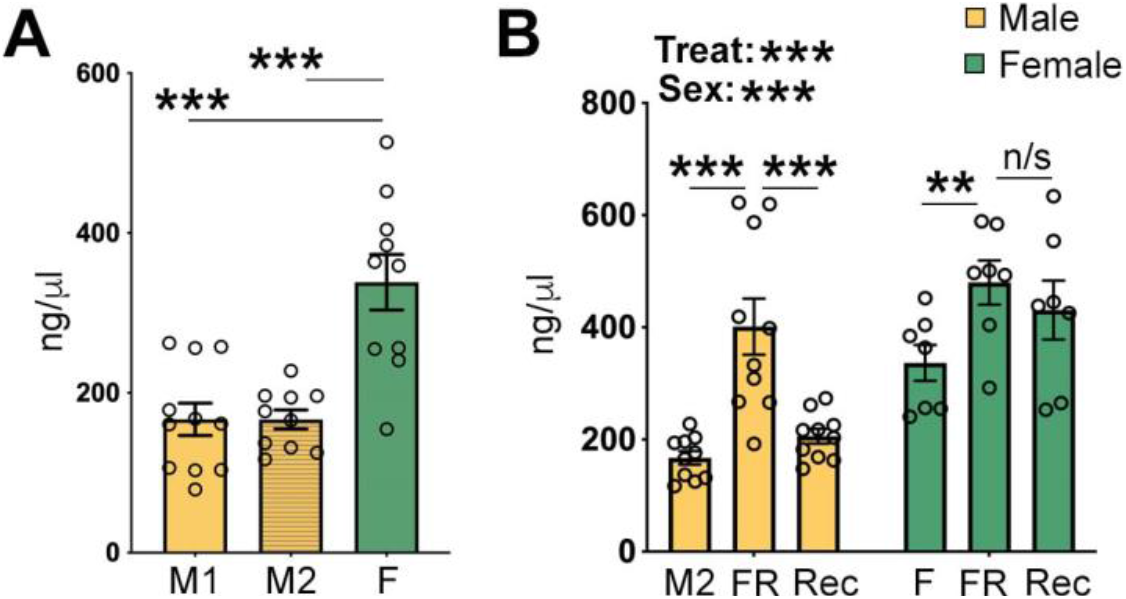
Plasma Corticosterone levels at baseline and with food restriction. (***A***) Female mice exhibited higher concentrations of CORT than males under baseline conditions. (**B**) CORT levels increased in both sexes during FR, but only females retained high levels once recovered from FR. Values are means ± *SEM*. Symbols are individual mouse data points. \**p* < 0.05, **<0.01, or ***<0.001

### 3.4 Digging discrimination in a model of autism and intellectual disability

Our interest in developing a more sensitive measure for digging behavior originated from the analysis of a mouse model of autism and intellectual disability, *Cc2d1a* conditional knock-out mice (cKO)(Oaks et al., 2017). *CC2D1A* loss of function leads to a spectrum of psychiatric presentations including severe to moderate intellectual disability, autism spectrum disorder, and aggressive behavior (Basel-Vanagaite et al., 2006; Loviglio et al., 2016; Manzini et al., 2014; Reuter et al., 2017). Mice where *Cc2d1a* is conditionally removed in the forebrain show an array of cognitive and social deficits, hyperactivity, and obsessive grooming, primarily found in males (Oaks et al., 2017; Zamarbide et al., 2019). *Cc2d1a* cKO males buried the same number of marbles as controls, but subsequent analysis of the videos identified a reduction in time spent digging (Oaks et al., 2017). We asked whether the digging discrimination test would be more sensitive in assessing changes in digging behavior.

We generated a cohort of male and female control (cre alone or homozygous floxed, Cont) and *Cc2d1a* cKO littermates and performed the DBD test (**Fig 4.** Cont M N=8, cKO M N=8, Cont F N=10, cKO F N=10). Despite a trend for females being more active, there was no significant difference in distance covered (**Fig. 4A**) and speed (**Fig. 4B**). In this transgenic strain, males of both genotypes performed significantly more digging than females when digging in the burrow and open area were added (**Fig. 4C**, Cont M: 230.6±24.9 s; cKO M: 226.4±34.7 s; Cont F: 106.3±23.8 s; cKO F: 115.6±27.7 s. Cont M/F p=0.017, cKO M/F p=0.04, 2-way ANOVA genotype p=0.982, sex=0.0002, **Suppl. Table 4** for additional statistics). However, digging efforts was differentially distributed to digging and burrowing. Digging in the open area showed a significant effect of both genotype and sex. *Cc2d1a* cKO males showed an increase in outside digging which was significantly higher than both groups of females and trended towards significance when compared to control males (**Fig. 4D**, Cont M: 39.6±12.8 s; cKO M: 84.7±19.3 s; Cont F: 19.5±5.1 s; cKO F: 25.6±7.5 s. Cont/cKO M p=0.076, cKO M/Cont F p=0.002, cKO M/cKO F p=0.006, 2-way ANOVA genotype p=0.03, sex p=0.002). As in the wild-type animals, this strain spent more time digging in the burrow than in the open area showing that burrowing is their preferred activity (**Fig. 4E**, Cont M: 0.81±0.06 s; cKO M: 0.55±0.11 s; Cont F: 0.61±0.12 s; cKO F: 0.62±0.12 s. **Suppl. Table 4**) and latencies to burrow were similar (**Suppl. Table 4**). Burrowing rates were only partially informative due to variability suggesting that cKO males were less efficient (**Suppl. Table 4**). As in the previous experiments, the percentage of material removed from the burrow was the most sensitive measure with a 62% reduction in burrowing and half the animals barely interacting with the substrate despite hovering in the vicinity of the burrow (**Fig. 4F,** WT M: 82.5±3.8%, cKO M: 31.5±16.2%, WT F: 49.7±12.8%, cKO F: 45.6±12.4%. M WT/cKO p=0.027, 2-way ANOVA genotype p=0.023, sex p=0.42).

**Figure 4.**
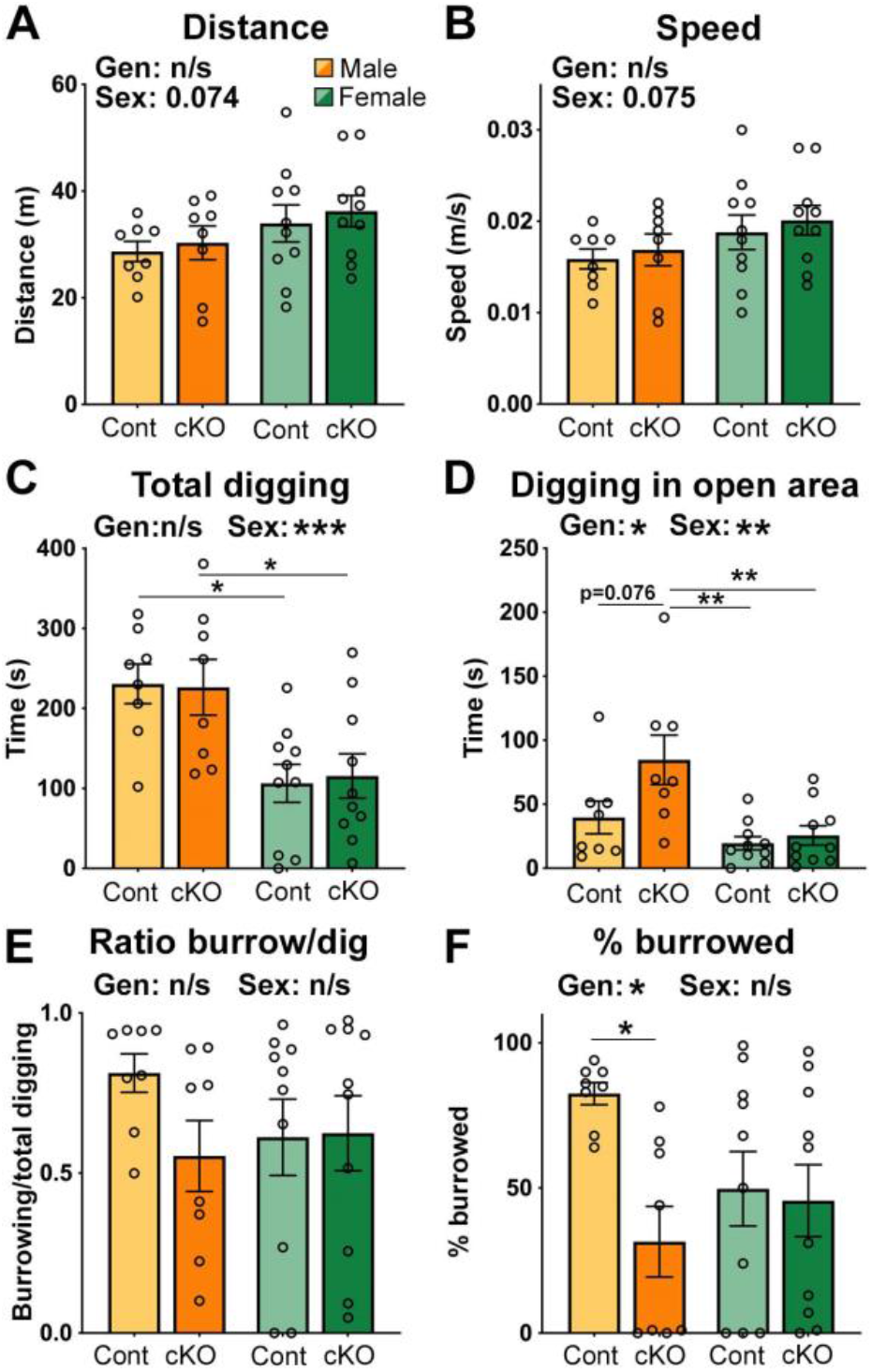
Cc2d1a cKO males show reduced burrowing performance. 4 cohorts of mice (Cont M: N=8; cKO M: N=8; Cont F: N=10; cKO F: N=10) were tested independently to assess the protocol sensitivity to an animal model of ASD. *(**A-B.**)* Between genotype male and females covered similar distances at equal speed, while females showed increased motor activity (**A.**) and speed (**B.**) when compared to males. (***C.***) Both control and cKO females showed significantly reduced total digging behavior when compared to males. (**D.**) cKO males spent significantly more time digging than both female cohorts and showed a trend towards more digging compared to control males. (**E.**) All animals spent more time burrowing than digging outside, though male cKOs and females showed great variability. (**F.**) cKO males burrowed significantly less material than wild type males, a difference not seen between female genotypes. Values are means ± *SEM*. Symbols are individual mouse data points. \**p* < 0.05, **p<0.01, or ***p<0.001 following multiple comparison tests. All unmarked comparisons were not significant. Additional statistical information is reported in Suppl. Table 4.

## Discussion

Measures of digging behavior are used to assess anxiety- and compulsive-like behaviors, and motor deficits in mice to study features of brain disorders (Bey & Jiang, 2014; Deacon et al., 2001; de Brower et al., 2019; Kazdoba et al., 2016; Thompson et al., 2019). However, many of the existing protocols are not able to elucidate digging motivation leading to inconsistencies in the interpretation of the experimental measures (Gyertyán, 1995; Njung’e & Handley, 1991; Thomas et al., 2009). In this study, we asked whether burrowing could be used in combination with exploratory digging for rapid assessment of motivation of digging behavior. We adapted a burrowing protocol developed by Deacon (2006a) that is sensitive to an array of motor and neurological deficits (Deacon, 2006a, 2012; Deacon et al., 2001, 2008) by adjusting the amount and texture of the burrowing substrate to produce measurable results in a shorter period of time.

We propose that in its simplest version the DBD test can be scored by using the percentage of material removed from the burrow as the burrowing measure and time digging in the open area as the exploratory digging measure. After exploring a variety of metrics obtained with both automated video-tracking and manual analysis, we found that weighing the material left inside the tube at the end of the test was the most sensitive measure of burrowing differences. While in the marble burying test, marbles can be buried and then unburied with vigorous digging leading to misleading results (Gyertyán, 1995), mice that remove soft bedding from the burrow do not push it back in. Exploratory digging must still be scored manually on video by trained raters unless there are appropriate algorithms that will identify specific digging posture and movement. When considering these two measures, the DBD protocol reliably identified multiple differences in burrowing and exploratory digging behavior in mice following an environmental change, i.e. food restriction, and a genetic mutation. In addition, this test revealed sex-specific changes in digging behavior that had not been observed in previous reports (Taylor et al., 2017).

Food restriction is known to alter foraging behavior and eating habits (Dell’Omo et al., 2000; Pankevich et al., 2010). While extended food restriction lasting over 10 days reduced overall activity and speed as previously shown (Tucci et al., 2006), total time spent digging was similar. However, male mice shifted towards spending more time digging in the open area and removed less material from the burrow. After *ad libitum* feeding was restored, digging and burrowing returned to baseline levels. In females, food restriction did not affect digging in the burrow or in the open area. While burrowing at baseline was not significantly different from males, females showed a much larger standard deviation and inconsistent performance in both the initial study cohort and control littermates for the *Cc2d1a* cKO. Interestingly, female burrowing performance improved to levels similar to males in the recovery trial after food restriction. Since burrowing has been shown to rely on both the hippocampus (Deacon & Rawlins, 2005) and frontal cortex (Deacon et al., 2003), it is possible that learning may contribute to better performance upon repetition of the test.

Mild extended food restriction induces a response in rodents in the hypothalamic-pituitary-adrenal (HPA) axis raising blood levels of CORT (Díaz-Muñoz et al., 2000; Méquinion et al., 2014; Scheurink et al., 1999; Yoshihara et al., 1996). In our studies, female mice showed higher CORT levels than both male cohorts at baseline as previously established (Kitay, 1961; Laviola et al., 2002). While CORT levels increased in both males and females with food restriction, they only returned to baseline in males. Female and male rodents have been shown by multiple groups to have distinct cellular and physiological responses and adaptation to stress and altered feeding regimens (Bale & Epperson, 2015; Massa & Correa, 2020; Rincón-Cortés et al., 2019). Correlation analysis showed that male digging behavior strongly correlated with CORT levels, but there was no correlation for females. While this CORT increase can be interpreted as a stress response, it is also thought to have an adaptive role leading to increased food anticipatory activity and recreational exercise (Díaz-Muñoz et al., 2000; Pankevich et al., 2010; Scheurink et al., 1999). It is possible that elevated CORT levels may be involved in increasing exploratory digging activity, but further studies will be needed to define how HPA axis activity affects digging in males.

Sex-specific digging changes were also observed in a mouse line deficient for the *Cc2d1a* gene, which is mutated in ID and ASD in humans. Removal of *Cc2d1a* in the cortex and hippocampus leads to hyperactivity and obsessive grooming in addition to cognitive and social deficits (Oaks et al., 2017; Yang et al., 2019). Reduced digging activity was identified in the marble burying test in *Cc2d1a* cKO males with no change in marble number (Oaks et al., 2017). The DBD test was more sensitive in defining digging changes with a substantial decrease in burrowing and a trend towards increased exploratory digging in male cKO mice. There was no difference between wild-type and female cKO mice. Females of any genotype dug less than mice in this strain indicating that despite a shared genetic background (C57BL/6N) there could also be baseline differences in digging due to husbandry and genetic manipulations. *Cc2d1a* cKOs have shown male-specific behavioral impairments in some behavioral tests linked to sex-specific signaling deficits in the hippocampus (Zamarbide et al., 2019), which may underlie the sex difference in these findings or compound with a different motivation for digging in males and females. Chen et al (2005) in studying the effects of senescence and aging on burrowing also showed that males and females differentially alter their burrowing performance with age and that this change may not be related to anxiety or novelty. One additional consideration is that corncob bedding commonly used in animal facilities has been shown to impact estrogen responses in rodents (Villalon Landeros et al., 2012) and to increase maternal care leading to reduced anxiety-like behaviors in the offspring (Sakhai et al., 2013). We used corncob because it was a familiar substrate, but different digging responses could be observed if animals are reared on other materials.

Overall, our studies show that digging is a complex and multidimensional behavior and that its motivation and performance must be explored in more detail, especially as it pertains to sex-specific changes. Our results reinforce the fact that mice are instinctually driven to dig and that digging choices are not random. Burrowing takes priority over exploration in both sexes, but males have a stronger drive to switch to exploratory digging than females. We cannot yet clearly assign a specific reason for this switch with the data at hand. It is possible that food restriction drives males to look for food outside the burrow and that food seeking is linked to CORT fluctuations and activity of the circuitry of the hypothalamus and reward pathways (Massa & Correa, 2020). A modified version of the DBD test where the free digging area is baited with food or where a food patch is provided may help to further define how mice choose between different digging modalities. Similarly, it is not known whether *Cc2d1a* cKO male mice lose their need to burrow due to increased anxiety or compulsion to dig outside. Additional studies could address the respective roles of the reward, fear, and motor circuitry in controlling digging motivation in this mouse strain and other strains carrying mutations linked to neurodevelopmental and neurological disorders. Finally, females used in our studies were sexually naïve, but different digging responses may appear when females are building a nest or protecting their young.

It is important to note that while we suggest focusing on percentage of material burrowed and time digging in open area as measures of burrowing and exploratory digging, the experimenter must always consider digging as a complex naturalistic behavior. We used a very conservative measure of digging, but additional motions to dig such as flicking substrate with one or both forefeet or wading into the substrate in a swim-like motion was observed by us and others (de Brouwer et al., 2019; Layne & Ehrhart, 1970). It would be interesting to explore these movements further in the future as they could be related to searching for food on the surface or disposing of dug soil. Spreading of the substrate removed from the burrow was an unexpected yet very consistent behavior which appears opposite to nest building behavior (Deacon, 2012; Neely et al., 2019). Burrowing behavior varies among rodents, so particular attention must be placed in understanding species-specific behavior (Reichman & Smith, 1990; Hu & Hoekstra, 2017; Metz et al., 2017). Wild house mice (*Mus musculus*) are known to seasonally clean their burrows of debris and spoiled food by pushing them out of the burrow (Schmid-Holmes et al., 2001). In addition, house mice usually have clear dirt paths or “runways” to the entry of their burrow systems (Avenant & Smith, 2003; Eriksson & Eldridge, 2014). This spreading behavior could reflect another motivated behavior linked to digging caused by an innate need to hide sediment from the excavation or clear the entrance to the burrow.

In closing, the current study underscores the need to consider digging behavior in laboratory mice as multifaceted and proposes a novel paradigm to probe digging motivation that can be completed with simple measures. Digging is tied to different aspects of a mouse well-being, from sheltering from dangers to obtaining and storing food, and like playing, it is a motor output integrating multiple circuits involved in learning and reward. The ability to distinguish between different types of digging in a single test may be beneficial to explore digging motivation and the underlying circuits.

## Acknowledgements

We are grateful to Adele Mossa and Pablo Munoz in the Manzini laboratory and Abigail Polter at the George Washington University for helpful discussions about the study. This research was supported by the National Institutes of Health NR01NS105000 and the Robert Wood Johnson Foundation.

## Author Contributions

All authors had full access to all the data in the study and take responsibility for the integrity of the data and the accuracy of the data analysis. *Conceptualization:* H.P.L, M.C.M; *Methodology:* H.P.L.; *Investigation:* H.P.L.; *Formal Analysis:* H.P.L, A.T.H, B.M.G, O.P.M., N.K.W., and M.C.M. *Writing – Original Draft:* A.T.H, B.M.G, and M.C.M; *Writing – Reviewing and Editing:* H.P.L., A.T.H, B.M.G, and M.C.M; *Visualization:* M.C.M.; *Supervision:* M.C.M.; *Funding Acquisition:* M.C.M.

## Data Accessibility

The data that support the findings are available from the corresponding author upon reasonable request.

## Supplementary Material

**Supplementary Table 1:** Values for baseline behavioral parameters and corticosterone levels shown in Figures 1 and 3. Values are presented ± s.e.m. Shapiro-Wilk test statistics are reported for the M1, M2 and F cohort at the bottom of each row and were used to determine whether the run parametric (one-way ANOVA) or non-parametric (Kruskal-Wallis) tests. The p values reported under the numbers refer to multiple comparison analyses (either Tukey’s for one-way ANOVA or Dunn’s for Kruskal-Wallis test).

|  | M1 N=11 | M2 N=13 | F N=10 |
| --- | --- | --- | --- |
| **Distance (m)**<br>**Fig.1B** | 33.72±2.41<br>p(M1/M2)=0.339 n/s<br>W=0.89/p=0.12 | 38.65±1.94<br>W=0.88/p=0.07 | 52.55±3.30<br>p(M1/F)=<0.0001 ***<br>p(M2/F)=0.0013 **<br>W=0.93/p=0.41 |
| **Speed (m/s)**<br>**Fig.1C** | 0.019±0.001<br>p(M1/M2)=0.339 n/s<br>W=0.88/p=0.21 | 0.021±0.001<br>W=0.87/p=0.06 | 0.029±0.002<br>p(M1/F)<0.0001 ***<br>p(M2/F)=0.0011 **<br>W=0.91/p=0.28 |
| **Burrow occupancy**<br>**(time in burrow/total)**<br>**Fig.1D** | 0.375±0.025<br>p(M1/M2)=0.988 n/s<br>W=0.96/p=0.81 | 0.380±0.017<br>W=0.89/p=0.09 | 0.353±0.024<br>p(M1/F)=0.768 n/s<br>p(M2/F)=0.663 n/s<br>W=0.90/p=0.23 |
| **Total digging (s)**<br>**Fig.1E** | 218.8±26.93<br>p(M1/M2)=0.185 n/s<br>W=0.99/p=0.996 | 151.3±14.15<br>W=0.92/p=0.22 | 169.6±36.90<br>p(M1/F)=0.206 n/s<br>p(M2/F)=0.999 n/s<br>W=0.72/p=0.002** |
| **Digging in open area (s)**<br>**Fig.1F** | 46.33±7.54<br>p(M1/M2)=0.783 n/s<br>W=0.94/p=0.54 | 32.14±4.74<br>W=0.94/p=0.56 | 64.63±27.72<br>p(M1/F) =0.702 n/s<br>p(M2/F) =0.292 n/s<br>W=0.60/p<0.0001*** |
| **Burrowing/total**<br>**digging ratio Fig.1G** | 0.741±0.089<br>p(M1/M2)=0.999 n/s<br>W=0.64/p=0.0003*** | 0.790±0.026<br>W=0.904/p=0.18 | 0.669±0.066<br>p(M1/F) =0.610 n/s<br>p(M2/F) =0.442 n/s<br>W=0.885/p=0.18 |
| **Latency to burrow (s)**<br>**Fig.1H** | 264.50±94.78<br>p(M1/M2)=0.266 n/s<br>W=0.58/p<0.0001 | 232.30±92.51<br>W=0.55/p<0.0001 | 228.90±52.31<br>p(M1/F)>0.999 n/s<br>p(M2/F)=0.283 n/s<br>W=0.86/p=0.07 |
| **Time to empty (s)**<br>**Fig.1I** | 1468±118.4<br>p(M1/M2)>0.999 n/s<br>W=0.78/p=0.005** | 1441.0±76.2<br>W=0.87/p=0.049* | 1543±67.1<br>p(M1/F) >0.999 n/s<br>p(M2/F)>0.999 n/s<br>W=0.91/p=0.29 |
| **Time burrowing (s)**<br>**Fig.1J** | 172.40±26.86<br>p(M1/M2)=0.094 n/s<br>W=0.93/p=0.49 | 119.2±11.78<br>W=0.88/p=0.08* | 105.0±13.37<br>p(M1/F) =0.040 *<br>p(M2/F)=0.832 n/s<br>W=0.91/p=0.30 |
| **Burrowing rate (g/min)**<br>**Fig.1K** | 4.22±0.78<br>p(M1/M2)=0.012 *<br>W=0.92/p=0.39 | 9.08±1.05<br>W=0.86/p=0.04* | 6.76±1.40<br>p(M1/F) =0.219 n/s<br>p(M2/F)=0.889 n/s<br>W=0.91/p=0.30 |
| **% material burrowed**<br>**Fig.1L** | 80.18±10.38<br>p(M1/M2)>0.999 n/s<br>W=0.62/p<0.0001*** | 94.00±2.16<br>W=0.78/p<0.0001*** | 69.50±10.73<br>p(M1/F) =0.301 n/s<br>p(M2/F)=0.069 n/s<br>W=0.82/p=0.03* |
| **Area of spread (% of cage area)**<br>**Fig. 1M** | 64.27±7.27<br>p(M1/M2)>0.999 n/s<br>W=0.84/p=0.03* | 74.00±2.55<br>W=0.97/p=0.92 | 57.00±6.61<br>p(M1/F) =0.499 n/s<br>p(M2/F)=0.057 n/s<br>W=0.87/p=0.11 |
| **Corticosterone**<br>**Fig.3A** | 166.90±20.22<br>p(M1/M2)>0.999 n/s<br>W=0.88/p=0.11 | 166.70±11.84<br>W=0.93/p=0.43 | 338.40±34.68<br>p(M1/F) <0.0001***<br>p(M2/F)<0.0001 ***<br>W=0.97/p=0.86 |

**Supplementary Table 2:**
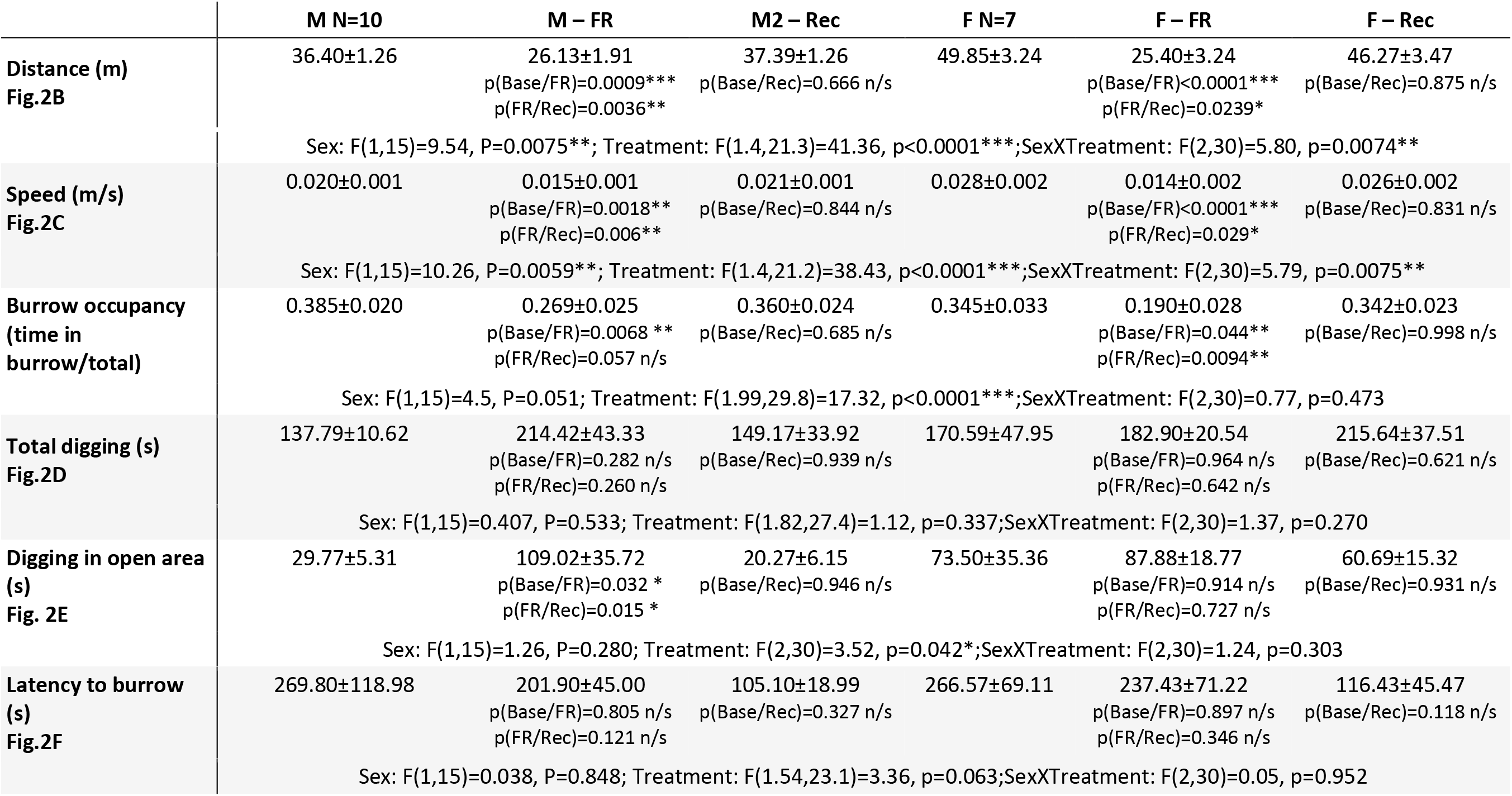
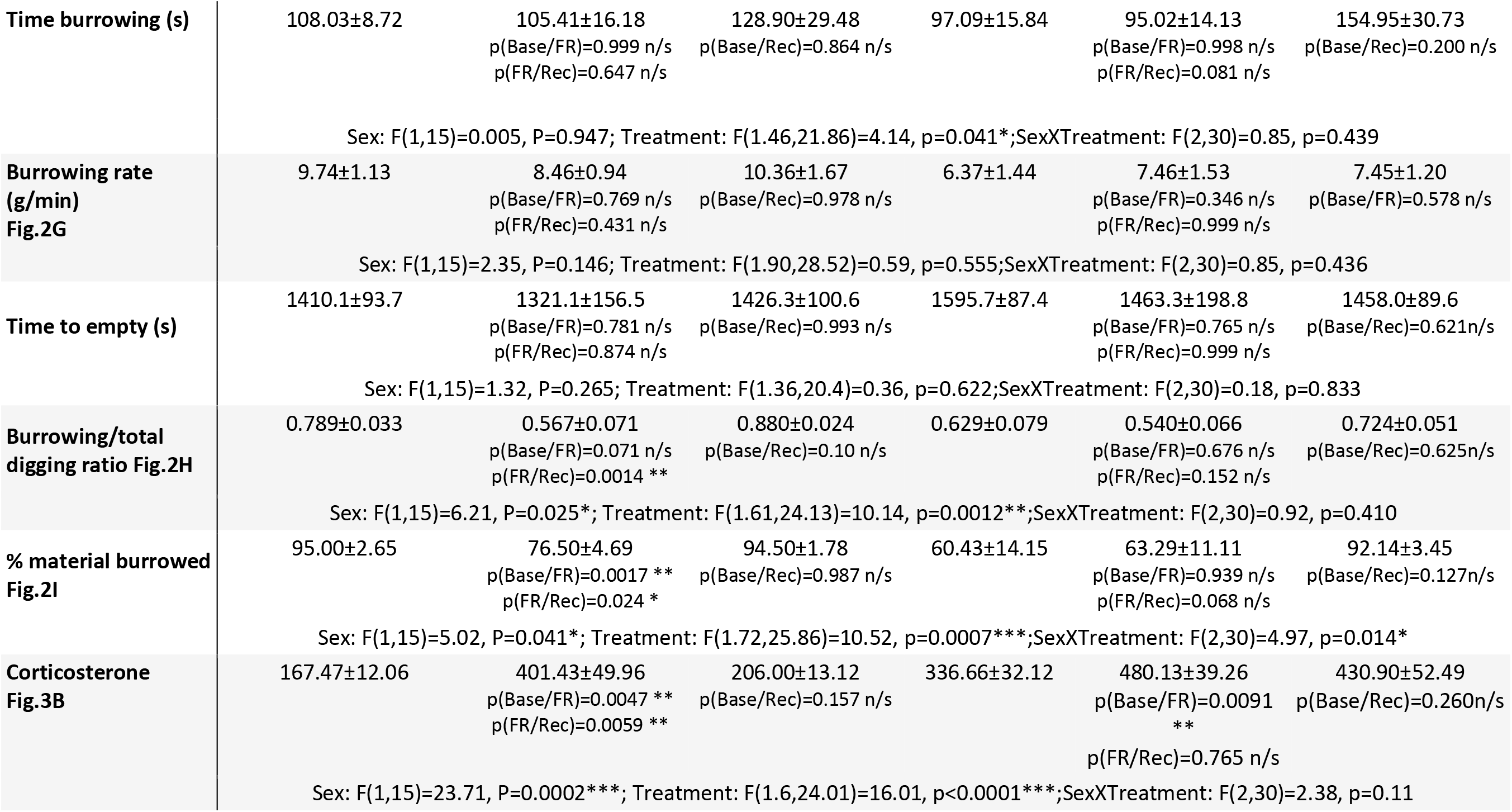
Values for behavioral parameters and corticosterone levels during the food restriction protocol shown in Figures 2 and 3 also including additional criteria mentioned in the Results. Values are presented ± s.e.m. P values for the baseline (Base), food restriction (FR) and recovery (Rec) studies are the result of Sidak multiple comparison tests following two-way repeated measures ANOVA testing for treatment and sex using a mixed-model to account for non-sphericity of data. ANOVA statistics for sex and treatment and sexXtreatment interaction are reported below each parameter.

**Supplementary Table 3:** Comparison of animals who escaped from food restriction trial with animals that completed the trial. Males that escaped appeared to be more active than the average in their cohort, but all mice that escaped showed burrowing parameters than the others with lower latency to burrow. Values are presented ± s.e.m. Results from food restriction cohort are below in parentheses. Parameters that could not be measured because the animals escaped from the arena are listed as not available (n/a).

|  | M2 - Base | M2 - FR | M2 - Rec | F - Base | F - FR | F - Rec |
| --- | --- | --- | --- | --- | --- | --- |
| **Distance (m)** | 46.18±6.07<br>(36.40±1.26) | n/a<br>(26.13±1.91) | 42.79±2.52<br>(37.39±1.26) | 54.43±7.39<br>(52.55±3.30) | n/a<br>(25.40±3.24) | 40.22±2.25<br>(46.27±3.47) |
| **Speed (m/s)** | 0.026±0.003<br>(0.020±0.001) | 0.013±0.003<br>(0.015±0.001) | 0.024±0.002<br>(0.021±0.001) | 0.030±0.004<br>(0.028±0.002) | 0.022±0.012<br>(0.014±0.002) | 0.022±0.001<br>(0.026±0.002) |
| **Latency to burrow (s)** | 107.33±1.45<br>(269.80±118.98) | 188.67±97.54<br>(201.90±45.00) | 99.00±42.33<br>(105.10±18.99) | 141.00±45.72<br>(266.57±69.11) | 72.00±29.70<br>(237.43±71.22) | 64.33±5.90<br>(116.43±45.47) |
| **Time in burrow (%)** | 36.1±3.5<br>(38.5±2.0) | n/a<br>(26.9±2.5) | 36.8±2.6<br>(36.0±2.4) | 37.23±3.0<br>(34.5±3.3) | n/a<br>(19.0±2.8) | 27.86±1.8<br>(34.2±2.3) |
| **Time to empty (s)** | 1545.3±110.5<br>(1410.1±93.7) | n/a<br>(1321.1±156.5) | 1258.0±201.2<br>(1426.3±100.6) | 1420.67±57.0<br>(1595.7±87.4) | n/a<br>(1463.3±198.8) | 1318.33±283.8<br>(1458.0±89.6) |
| **% material burrowed** | 90.80±2.98<br>(95.00±2.65) | n/a<br>(76.50±4.69) | 95.41±2.96<br>(94.50±1.78) | 90.69±2.23<br>(60.43±14.15) | n/a<br>(63.29±11.11) | 94.76±2.66<br>(92.14±3.45) |
| **Corticosterone** | 198.67±8.50<br>(167.47±12.06) | 389.60±79.21<br>(401.43±49.96) | 120.29±24.13<br>(206.00±13.12) | 342.40±103.97<br>(336.66±32.12) | 514.03±181.06<br>(480.13±39.26) | 575.13±49.21<br>(430.90±52.49) |

Abbreviations: FR=food restriction; Rec=recovery; n/a= not available

**Supplementary Table 4:**
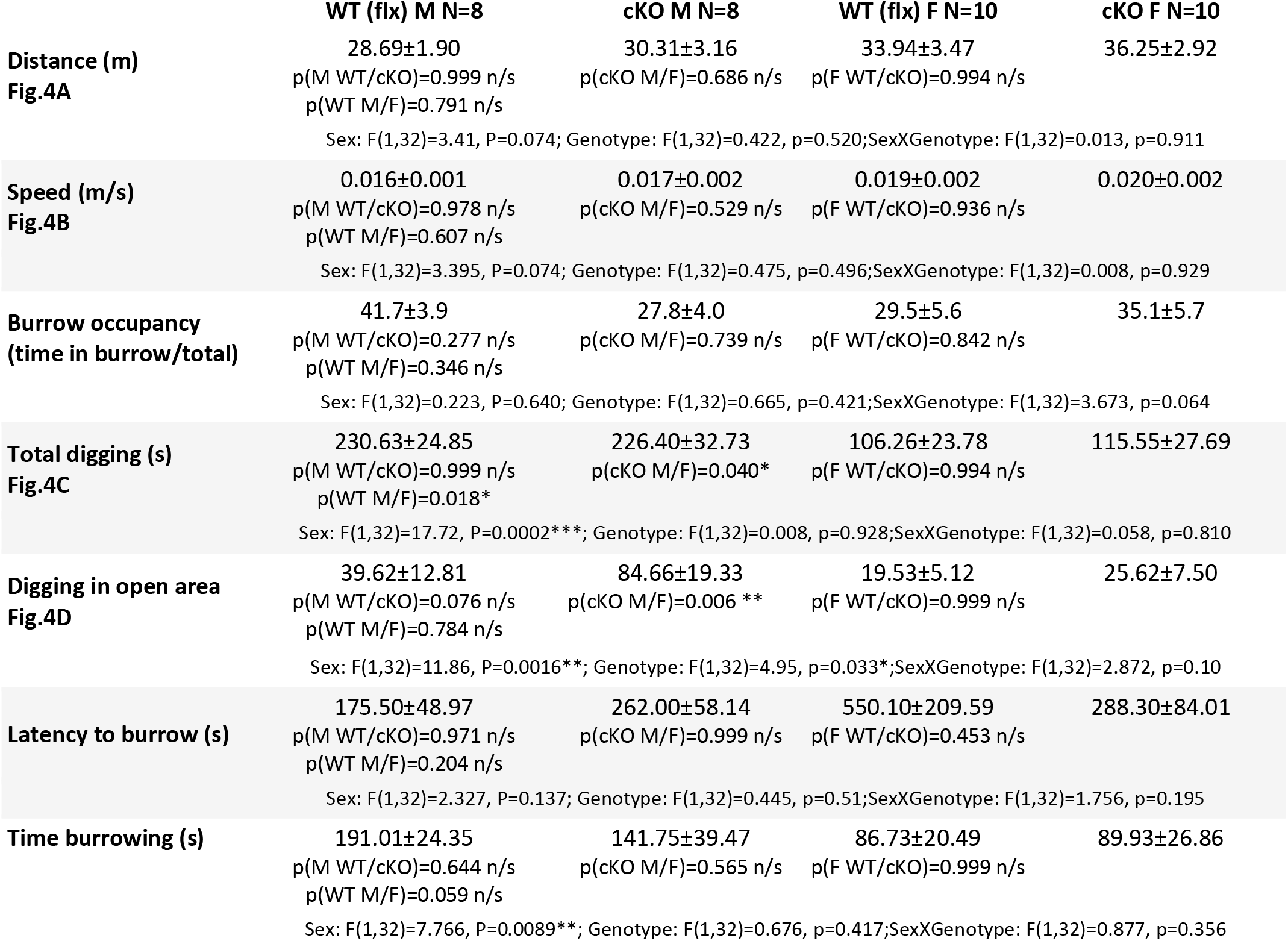
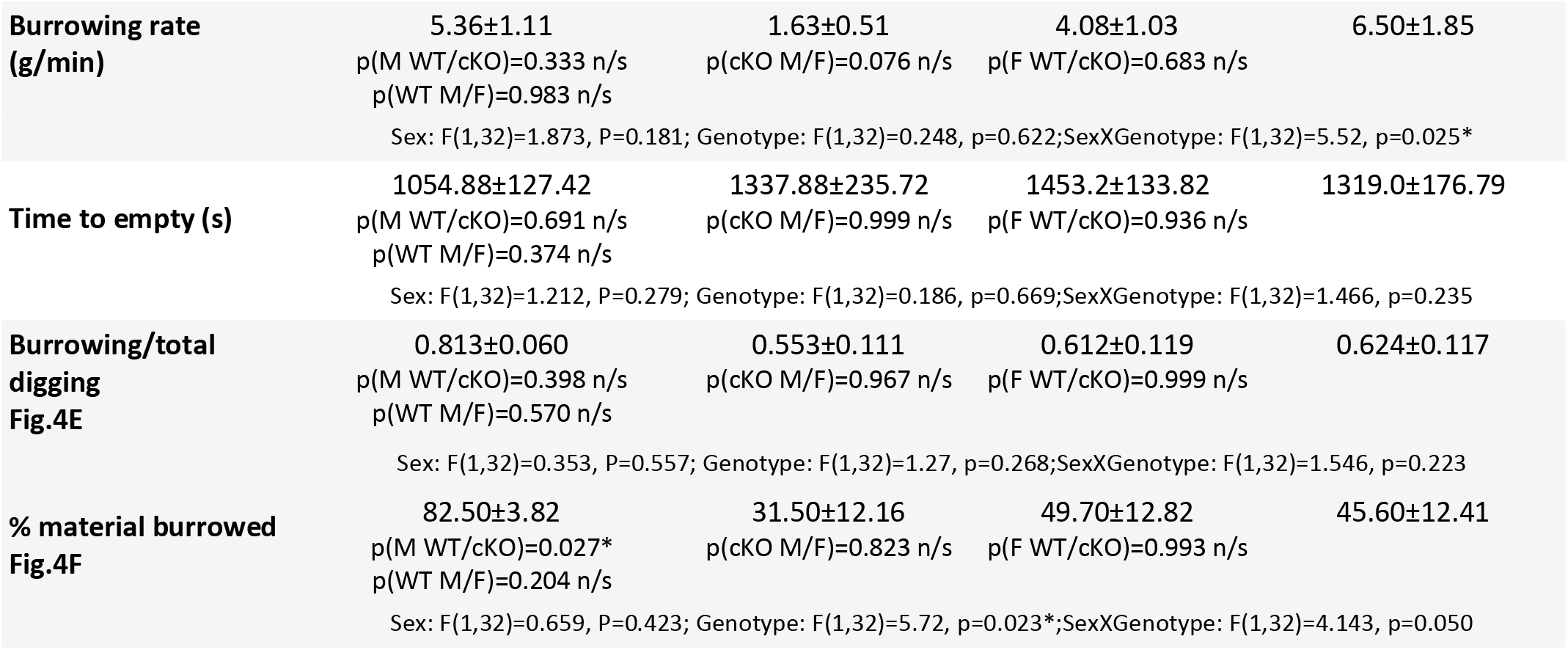
Values for behavioral parameters for the Cc2d1a cKO mice reported in Figure 4. Values are presented ± s.e.m. P values are the result of Sidak multiple comparison tests following two-way ordinary ANOVA testing for genotype and sex. ANOVA statistics for sex and treatment and sexXtreatment interaction are reported below each parameter.

## References

American Psychiatric Association. (2013). Diagnostic and Statistical Manual of Mental Disorders (Fifth Edition). American Psychiatric Association.

Arakawa, H., Blanchard, D. C., & Blanchard, R. J. (2007). Colony formation of C57BL/6J mice in visible burrow system: Identification of eusocial behaviors in a background strain for genetic animal models of autism. Behavioural Brain Research, 176(1), 27–39.

Avenant, N. L., & Smith, V. R. (2003). The microenvironment of house mice on Marion Island (sub-Antarctic). Polar Biology, 26(2), 129–141.

Bakdash, J.Z., & Marusich, L. R. (2019). Repeated measures correlation. Frontiers in Psychology, 8:456 doi: 10.3389/fpryg.2017.00456

Bale, T. L., & Epperson, C. N. (2015). Sex differences and stress across the lifespan. Nature Neuroscience, 18(10), 1413–1420.

Basel-Vanagaite, L., Attia, R., Yahav, M., Ferland, R. J., Anteki, L., Walsh, C. A., Olender, T., Straussberg, R., Magal, N., Taub, E., Drasinover, V., Alkelai, A., Bercovich, D., Rechavi, G., Simon, A. J., & Shohat, M. (2006). The CC2D1A, a member of a new gene family with C2 domains, is involved in autosomal recessive non‐syndromic mental retardation. Journal of Medical Genetics, 43(3), 203–210.

Bey, A. L., & Jiang, Y. (2014). Overview of mouse models of autism spectrum disorders. Current Protocols in Pharmacology, 66, 5.66.1–26.

Broekkamp, C. L., Rijk, H. W., Joly-Gelouin, D., & Lloyd, K. L. (1986). Major tranquillizers can be distinguished from minor tranquillizers on the basis of effects on marble burying and swim-induced grooming in mice. European Journal of Pharmacology, 126(3), 223–229.

Bruins Slot, L. A., Bardin, L., Auclair, A. L., Depoortere, R., & Newman-Tancredi, A. (2008). Effects of antipsychotics and reference monoaminergic ligands on marble burying behavior in mice. Behavioural Pharmacology, 19(2), 145–152.

de Brouwer, G., Fick, A., Harvey, B. H., & Wolmarans, D. W. (2019). A critical inquiry into marble-burying as a preclinical screening paradigm of relevance for anxiety and obsessive-compulsive disorder: Mapping the way forward. Cognitive, Affective & Behavioral Neuroscience, 19(1), 1–39.

Deacon, R. M. (2006a). Burrowing in rodents: A sensitive method for detecting behavioral dysfunction. Nature Protocols, 1(1), 118–121.

Deacon, R. M. (2006b). Digging and marble burying in mice: Simple methods for in vivo identification of biological impacts. Nature Protocols, 1(1), 122–124.

Deacon, R. M. (2012). Assessing Burrowing, Nest Construction, and Hoarding in Mice. JoVE (Journal of Visualized Experiments), 59, e2607.

Deacon, R. M., Cholerton, L. L., Talbot, K., Nair-Roberts, R. G., Sanderson, D. J., Romberg, C., Koros, E., Bornemann, K. D., & Rawlins, J. N. P. (2008). Age-dependent and - independent behavioral deficits in Tg2576 mice. Behavioural Brain Research, 189(1), 126–138.

Deacon, R. M., Penny, C., & Rawlins, J. N. P. (2003). Effects of medial prefrontal cortex cytotoxic lesions in mice. Behavioural Brain Research, 139(1–2), 139–155.

Deacon, R. M., Raley, J. M., Perry, V. H., & Rawlins, J. N. (2001). Burrowing into prion disease. Neuroreport, 12(9), 2053–2057.

Deacon, R. M., & Rawlins, J. N. P. (2005). Hippocampal lesions, species-typical behaviours and anxiety in mice. Behavioural Brain Research, 156(2), 241–249.

Dell’Omo, G., Ricceri, L., Wolfer, D. P., Poletaeva, I. I., & Lipp, H.-P. (2000). Temporal and spatial adaptation to food restriction in mice under naturalistic conditions. Behavioural Brain Research, 115(1), 1–8.

Díaz-Muñoz, M., Vázquez-Martínez, O., Aguilar-Roblero, R., & Escobar, C. (2000). Anticipatory changes in liver metabolism and entrainment of insulin, glucagon, and corticosterone in food-restricted rats. American Journal of Physiology. Regulatory, Integrative and Comparative Physiology, 279(6), R2048–2056.

Dudek, B. C., Adams, N., Boice, R., & Abbott, M. E. (1983). Genetic influences on digging behaviors in mice (Mus musculus) in laboratory and seminatural settings. Journal of Comparative Psychology, 97(3), 249–259.

Eriksson, B., & Eldridge, D. (2014). Surface destabilisation by the invasive burrowing engineer Mus musculus on a sub-Antarctic island. Geomorphology, 223, 61–66.

Guarnieri, D. J., Brayton, C. E., Richards, S. M., Maldonado-Aviles, J., Trinko, J. R., Nelson, J., Taylor, J. R., Gourley, S. L., & DiLeone, R. J. (2012). Gene profiling reveals a role for stress hormones in the molecular and behavioral response to food restriction. Biological Psychiatry, 71(4), 358–365.

Gyertyán, I. (1995). Analysis of the marble burying response: Marbles serve to measure digging rather than evoke burying. Behavioural Pharmacology, 6(1), 24–31.

Hayashi, E., Kuratani, K., Kinoshita, M., & Hara, H. (2010). Pharmacologically distinctive behaviors other than burying marbles during the marble burying test in mice. Pharmacology, 86(5–6), 293–296.

Hörndli, C., Wong, E., Ferris, E., Bennett, K., Steinwand, S., Rhodes, A., Fletcher, P., & Gregg, C. (2019). Complex Economic Behavior Patterns Are Constructed from Finite, Genetically Controlled Modules of Behavior. Cell Reports, 28, 1814–1829.e6.

Hu, C. K., Hoekstra, H. E. (2017). Peromyscus burrowing: A model system for behavioral evolution. Seminars in Cell & Developmental Biology, 61, 107–114.

Jirkof, P. (2014). Burrowing and nest building behavior as indicators of well-being in mice. Journal of Neuroscience Methods, 234, 139–146.

Jirkof, P., Cesarovic, N., Rettich, A., Nicholls, F., Seifert, B., & Arras, M. (2010). Burrowing Behavior as an Indicator of Post-Laparotomy Pain in Mice. Frontiers in Behavioral Neuroscience, 4, 165.

Kazdoba, T. M., Leach, P. T., Yang, M., Silverman, J. L., Solomon, M., & Crawley, J. N. (2016). Translational Mouse Models of Autism: Advancing Toward Pharmacological Therapeutics. Current Topics in Behavioral Neurosciences, 28, 1–52.

Kitay, J. I. (1961). Sex differences in adrenal cortical secretion in the rat. Endocrinology, 68, 818–824.

Latham, N., & Mason, G. (2004). From house mouse to mouse house: The behavioural biology of free-living Mus musculus and its implications in the laboratory. Applied Animal Behaviour Science, 86(3), 261–289.

Laviola, G., Adriani, W., Morley-Fletcher, S., & Terranova, M. L. (2002). Peculiar response of adolescent mice to acute and chronic stress and to amphetamine: Evidence of sex differences. Behavioural Brain Research, 130(1–2), 117–125.

Layne, J. N., & Ehrhart, L. M. (1970). Digging behavior of four species of deer mice (Peromyscus). American Museum Novitates, 2429, 1–16

Loviglio, M. N., Beck, C. R., White, J. J., Leleu, M., Harel, T., Guex, N., Niknejad, A., Bi, W., Chen, E. S., Crespo, I., Yan, J., Charng, W.-L., Gu, S., Fang, P., Coban-Akdemir, Z., Shaw, C. A., Jhangiani, S. N., Muzny, D. M., Gibbs, R. A., … Reymond, A. (2016). Identification of a RAI1-associated disease network through integration of exome sequencing, transcriptomics, and 3D genomics. Genome Medicine, 8, 105.

Luigjes, J., Lorenzetti, V., de Haan, S., Youssef, G. J., Murawski, C., Sjoerds, Z., van den Brink, W., Denys, D., Fontenelle, L. F., & Yücel, M. (2019). Defining Compulsive Behavior. Neuropsychology Review, 29(1), 4–13.

Manzini, M. C., Xiong, L., Shaheen, R., Tambunan, D. E., Di Costanzo, S., Mitisalis, V., Tischfield, D. J., Cinquino, A., Ghaziuddin, M., Christian, M., Jiang, Q., Laurent, S., Nanjiani, Z. A., Rasheed, S., Hill, R. S., Lizarraga, S. B., Gleason, D., Sabbagh, D., Salih, M. A., … Walsh, C. A. (2014). CC2D1A Regulates Human Intellectual and Social Function as well as NF-κB Signaling Homeostasis. Cell Reports, 8(3), 647–655.

Massa, M. G., & Correa, S. M. (2020). Sexes on the brain: Sex as multiple biological variables in the neuronal control of feeding. Biochimica et Biophysica Acta (BBA) - Molecular Basis of Disease, 1866(10), 165840.

Méquinion, M., Caron, E., Zgheib, S., Stievenard, A., Zizzari, P., Tolle, V., Cortet, B., Lucas, S., Prévot, V., Chauveau, C., & Viltart, O. (2014). Physical activity: Benefit or weakness in metabolic adaptations in a mouse model of chronic food restriction? American Journal of Physiology-Endocrinology and Metabolism, 308(3), E241–E255.

Metz, H. C., Bedford, N. L., Pan, Y. L., & Hoekstra, H. E. (2017). Evolution and Genetics of Precocious Burrowing Behavior in Peromyscus Mice. Current Biology: CB, 27(24), 3837–3845.e3.

Neely, C. L. C., Pedemonte, K. A., Boggs, K. N., & Flinn, J. M. (2019). Nest Building Behavior as an Early Indicator of Behavioral Deficits in Mice. JoVE (Journal of Visualized Experiments), 152, e60139.

Njung’e, K., & Handley, S. L. (1991). Evaluation of marble-burying behavior as a model of anxiety. Pharmacology, Biochemistry, and Behavior, 38(1), 63–67.

Oaks, A. W., Zamarbide, M., Tambunan, D. E., Santini, E., Di Costanzo, S., Pond, H. L., Johnson, M. W., Lin, J., Gonzalez, D. M., Boehler, J. F., Wu, G. K., Klann, E., Walsh, C. A., & Manzini, M. C. (2017). Cc2d1a Loss of Function Disrupts Functional and Morphological Development in Forebrain Neurons Leading to Cognitive and Social Deficits. Cerebral Cortex, 27(2), 1670–1685.

Palanza, P., Morley-Fletcher, S., & Laviola, G. (2001). Novelty seeking in periadolescent mice: Sex differences and influence of intrauterine position. Physiology & Behavior, 72(1–2), 255–262.

Pankevich, D. E., Teegarden, S. L., Hedin, A. D., Jensen, C. L., & Bale, T. L. (2010). Caloric restriction experience reprograms stress and orexigenic pathways and promotes binge eating. Journal of Neuroscience, 30(48), 16399–16407.

Powell, F., & Banks, P. (2004). Do house mice modify their foraging behaviour in response to predator odours and habitat? Animal Behaviour, 67, 753–759

Reichman, O. J., & Smith S. C. (1990) Burrows and burrowing behavior by mammals. In H.H. Genoways ed., Current Mammalogy, Plenum Press, 197–244

Reuter, M. S., Tawamie, H., Buchert, R., Hosny Gebril, O., Froukh, T., Thiel, C., Uebe, S., Ekici, A. B., Krumbiegel, M., Zweier, C., Hoyer, J., Eberlein, K., Bauer, J., Scheller, U., Strom, T. M., Hoffjan, S., Abdelraouf, E. R., Meguid, N. A., Abboud, A., … Abou Jamra, R. (2017). Diagnostic Yield and Novel Candidate Genes by Exome Sequencing in 152 Consanguineous Families With Neurodevelopmental Disorders. JAMA Psychiatry, 74(3), 293–299.

Rincón-Cortés, M., Herman, J. P., Lupien, S., Maguire, J., & Shansky, R. M. (2019). Stress: Influence of sex, reproductive status and gender. Neurobiology of Stress, 10, 100155.

Sakhai, S.A., Preslik, J., Francis, D.D. Influence of housing variables on the development of stress-sensitive behaviors in the rat. Physiology & Behavior, 120:156–163

Scheurink, A. J., Ammar, A. A., Benthem, B., van Dijk, G., & Södersten, P. A. (1999). Exercise and the regulation of energy intake. International Journal of Obesity and Related Metabolic Disorders: Journal of the International Association for the Study of Obesity, 23 Suppl 3, S1–6.

Schmid-Holmes, S., Drickamer, L. C., Robinson, A. S., & Gillie, L. L. (2001). Burrows and Burrow-Cleaning Behavior of House Mice (Mus Musculus Domesticus). The American Midland Naturalist, 146(1), 53–62.

Sherwin, C. M., Haug, E., Terkelsen, N., & Vadgama, M. (2004). Studies on the motivation for burrowing by laboratory mice. Applied Animal Behaviour Science, 88(3), 343–358.

Sluyter, F., Korte, S. M., Bohus, B., & Van Oortmerssen, G. A. (1996). Behavioral stress response of genetically selected aggressive and nonaggressive wild house mice in the shock-probe/defensive burying test. Pharmacology, Biochemistry, and Behavior, 54(1), 113–116.

Taylor, G. T., Lerch, S., & Chourbaji, S. (2017). Marble burying as compulsive behaviors in male and female mice. Acta Neurobiologiae Experimentalis, 77(3), 254–260.

Thomas, A., Burant, A., Bui, N., Graham, D., Yuva-Paylor, L. A., & Paylor, R. (2009). Marble burying reflects a repetitive and perseverative behavior more than novelty-induced anxiety. Psychopharmacology, 204(2), 361–373.

Thompson, S. L., Welch, A. C., Ho, E. V., Bessa, J. M., Portugal-Nunes, C., Morais, M., Young, J. W., Knowles, J. A., & Dulawa, S. C. (2019). Btbd3 expression regulates compulsive-like and exploratory behaviors in mice. Translational Psychiatry, 9(1), 1–14.

Tsien, J. Z., Chen, D. F., Gerber, D., Tom, C., Mercer, E. H., Anderson, D. J., Mayford, M., Kandel, E. R., & Tonegawa, S. (1996). Subregion- and cell type-restricted gene knockout in mouse brain. Cell, 87(7), 1317–1326.

Tucci, V., Hardy, A., & Nolan, P. M. (2006). A comparison of physiological and behavioural parameters in C57BL/6J mice undergoing food or water restriction regimes. Behavioural Brain Research, 173(1), 22–29.

Villalon Landeros, R., Morisseau, C., Yoo, H.J., Fu, S.H., Hammock, B.D., Trainor, B.C. (2012) Corncob bedding alters the effects of estrogens on aggressive behavior and reduces estrogen receptor-alpha expression in the brain. Endocrinology, 153(2):949–953

Webster, D. G., Williams, M. H., Owens, R. D., Geiger, V. B., & Dewsbury, D. A. (1981). Digging behavior in 12 taxa of muroid rodents. Animal Learning & Behavior, 9(2), 173–177.

Yang, C.-Y., Yu, T.-H., Wen, W.-L., Ling, P., & Hsu, K.-S. (2019). Conditional Deletion of CC2D1A Reduces Hippocampal Synaptic Plasticity and Impairs Cognitive Function through Rac1 Hyperactivation. Journal of Neuroscience, 39(25), 4959–4975.

Yoshihara, T., Honma, S., Katsuno, Y., & Honma, K. (1996). Dissociation of paraventricular NPY release and plasma corticosterone levels in rats under food deprivation. The American Journal of Physiology, 271(2 Pt 1), E239–245.

Zamarbide, M., Mossa, A., Muñoz-Llancao, P., Wilkinson, M. K., Pond, H. L., Oaks, A. W., & Manzini, M. C. (2019). Male-Specific cAMP Signaling in the Hippocampus Controls Spatial Memory Deficits in a Mouse Model of Autism and Intellectual Disability. Biological Psychiatry, 85(9), 760–768.

